# Clinically oriented system-based embryology; a significant course in clinical practice

**DOI:** 10.1101/2020.02.13.946988

**Authors:** Samira Moghadam, Hamid Gharebaghi, Mohamad Javad Mohamadi, Ahmad Farokhi, Mehrdad Ghorbanlou, Mitra Arianmanesh

## Abstract

**Introduction:** Embryology is a branch of medical sciences in developmental biology. Since the knowledge of embryology is of special significance for medical students, this study was conducted with the purpose of elucidating the viewpoint of medical students of Zanjan University of Medical Sciences in Iran on the application of embryology courses in fulfilling clinical purposes.

**Methods:** This cross-sectional study was conducted in 2018-2019 with a census method on all clinical medical students (trainees and interns). To collect medical students’ opinions, the researcher-designed questionnaire was used. The validity of the questionnaire was confirmed by embryology experts by content validity ratio (CVR) and factor analysis and the reliability of the questionnaire Cronbach’s alpha coefficient (0.9), respectively. Data were analyzed by SPSS (version 24) and Lisrel software. Values with P<0.05 were considered significant.

**Results:** Descriptive statistics in the field of general and system-based embryology demonstrated that the topics of birth defects and development of body cavities (diaphragm development) have the most influence on clinical practice of medical students.

**Conclusions:** It seems that more focus on system-based embryology courses leads to a better performance of medical students in clinical courses.

## Introduction

Embryology is a branch of biological sciences in which students learn how the embryo or fetus develops from a single cell to a newborn in 9 months; the study of these complex phenomena is called embryology [1]. Like other medical sciences, embryology has been developed over the years. This fact brings up the challenge of how to present the course in medical schools [2]. To better correlate the basic sciences and clinical practice, studies suggest advance organizers, teaching through a clinical scenario, and students’ preparation before the theoretical course [3, 4]. In planning a medical curriculum, it is necessary to contain courses which are of value for training future physicians; embryology is one of those necessary courses assisting under-trained physicians to understand how our body becomes a human being, perceive birth defects and teratology as common topics which a physician encounters, and provide a logical basis for understanding the overall organization of the human body [5].

Embryology has had problems with acquiring an appropriate position in the medical curriculum. Neglecting embryology as a significant course may be due to several reasons including students’ thought toward embryology in which they pay more attention to anatomy rather than embryology or the lack of providing clinical information when teaching the course [6, 7].

To elucidate the importance of embryology course in clinical performance, a researcher-designed questionnaire was used to evaluate the effects of embryology course on clinical concepts of disease by asking the clinical medical students’ (trainees and interns) to fill the questioner.

## Methods

### Method of sampling

In this cross-sectional study performed in 2018-2019, because of our universe was limited to about 100 clinical medical students (trainees and interns) at the time of the study at the Zanjan University of Medical Sciences, therefore, we don’t have any sampling, and we use the census method for all the 100 clinical medical students. This study was confirmed in the ethics committee of Zanjan University of Medical Sciences with an ethics code of ZUMS.REC.1395.191.

### Designing a questionnaire

The research-design multiple-choice questionnaire was used to collect data. It consisted of two parts: the first part included questions about the application of embryology course in training and internship periods, and the second part included demographic information. The first part included 25 questions consisting of subcategories of general embryology (5 questions), system-based embryology (9 questions), teaching method (4 questions), educational assistance tools (3 questions), and clinical performance of medical students (4 questions). To maximize the anonymity of the respondents, demographic questions including gender, age, and education level were asked at the end of the questionnaire. A Likert scale consisting of five levels (1: strongly disagree, 2: disagree, 3: neither agree nor disagree, 4: agree, and 5: strongly agree) was applied in this study. In purpose Pre-testing questionnaires and determining the strengths and weaknesses of the questionnaire like, question format, wording, and order and for making sure that everyone in our main population can complete the questionnaire we find 5-10 people who were too close to our target group to pretest it. To determine the reliability and validity, 50 questionnaires were distributed among the medical students approaching the end of their education in Zanjan University of medical sciences. The validity of the questionnaire was confirmed by embryology experts by content validity ratio (CVR) and factor analysis and the reliability of the questionnaire Cronbach’s alpha coefficient (0.9), respectively (Table 1).

**Table 1:**
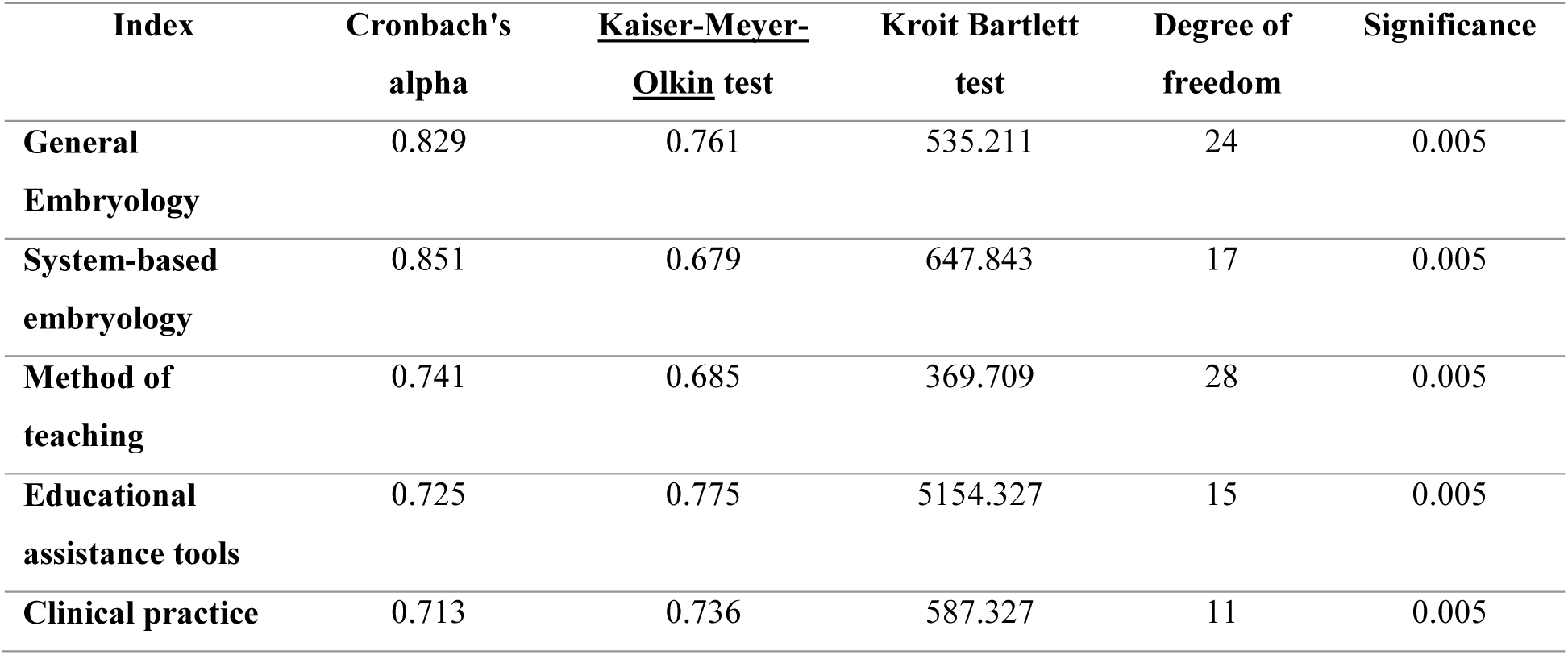
Validity and reliability test of the questionnaire by Cronbach’s alpha and factor analysis

### Statistical Analysis

The main analytical approach in this study was structural equation modeling. To apply this approach, at first, some missing data were applied by maximum-likelihood based missing data method and expectation maximization technique. Statistical analysis was performed using SPSS software (Version 24). To assure the normal distribution of data which is a basic condition to apply parametric tests, data were analyzed by Anderson–Darling test. In this study, variants of general embryology, system-based embryology, and clinical performance were normally distributed. For the educational assistance tools in embryology (P=0.04) type conversion was applied. The type conversion method applied was Johnson’s conversion method. To test the hypotheses and the impact of the independent variants, structural equation modeling was used in Lisrel software. In this method, the path coefficient within the critical value (−1.96 >T> 1.96) was considered significant.

## Results

### Demographic data

Data indicated 56% of participants were female and 44% were male. Also, 86% of participants were trainees and 14% were interns. Among the participants, 57% were between 23 to 25, 35% under 23 and 8% above 25 years old.

### Hypothesis testing

Figure 1 is the final model of this study which is designed according to the theoretical principles of the study. T-values are used to confirm or reject the study hypotheses. Figure 2 is the standard form of the assumed model. In this model, comparison and ranking of the variants are possible in which the higher path coefficient expresses the higher influence. Positive values express the positive effect which increases the amount of dependent variable, and negative values express a negative effect which decreases the amount of dependent variable.

**Figure 1.**
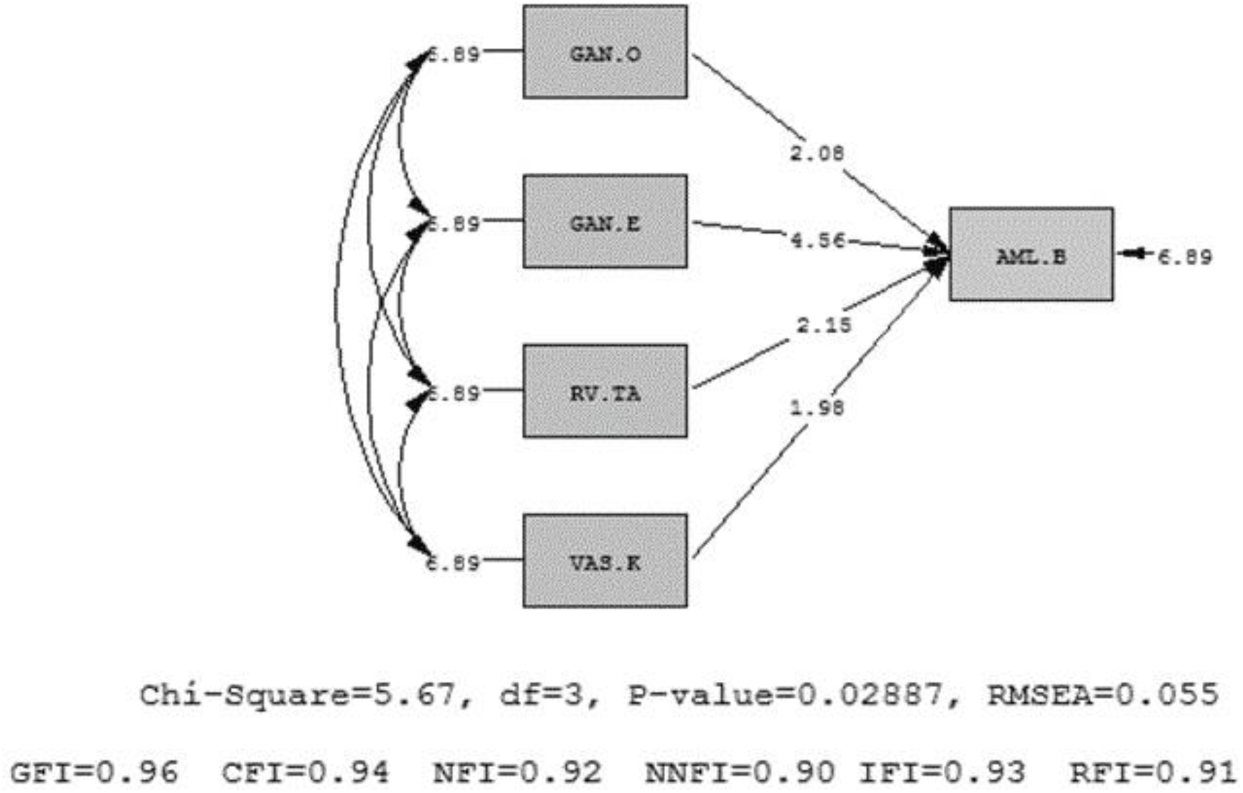
Final structural model test. In this case, it is possible to compare and rank the variables. The high value of path coefficient means higher impact, a positive value means a positive impact, and a negative value means a negative impact.

**Figure 2.**
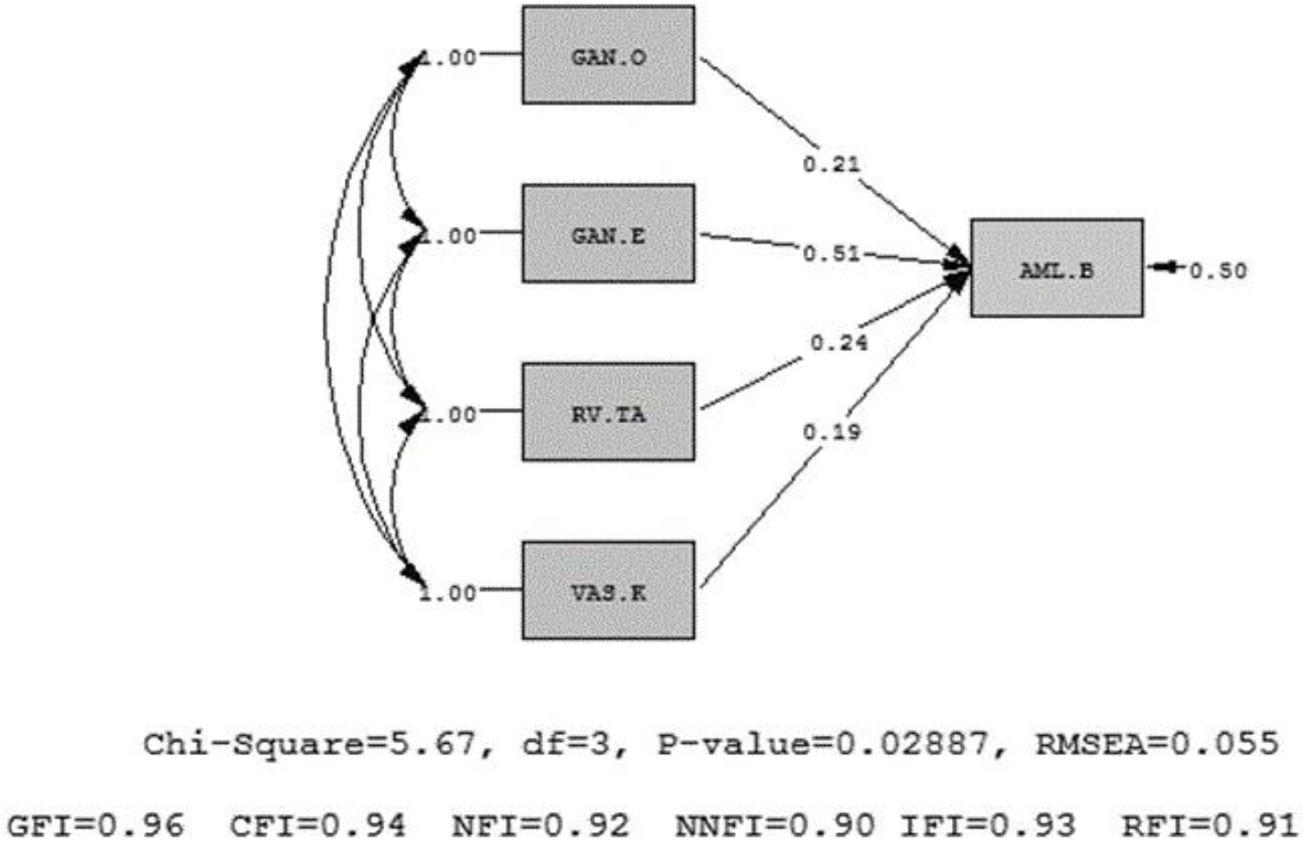
Final structural model test in T-Values mode. It is developed according to the conceptual model of the research and supported by theoretical foundations. In this case, it can be confirmed or rejected by research hypotheses.

Using structural equation modeling with LISREL 8.7 [8], hypotheses were tested. The overall fit of this model was good. The chi-square value on degrees of freedom df / χ2 is the first criteria to judge the fitness of the model. This criterion is used for single dimensional structures and its value must always be less than 3. The value of this index in this model is 1.92 which is smaller than 3. The root mean square error of approximation (RMSEA) is 0.055 which is lower than the maximum allowed amount of 0.08. According to figure 2, Goodness of Fit Index (GFI: 0.96), Confirmatory Fit Index (CFI: 0.94), Normal Fit Index (NFI: 0.92), Non-Normed Fit Index (NNFI: 0.90), Increasing Fitness Index (IFI: 0.93), and Relative Fitness Index (RFI: 0.92) are all on threshold of fitness allowed values. Therefore, hypotheses can be tested with assurance.

### The effect of general embryology on the clinical performance of medical students

According to the pathways analysis pattern and table 2, the standard pathway coefficient of general embryology with respect to the clinical performance of medical students was 0.21 (Figure 2). Therefore, regarding the t value of this pathway (t= 2.08 > 1.96) (Figure 1), it can be concluded that general embryology affects the clinical practice of medical students.

**Table 2:** Structural equation model.

| Variants | Standard coefficients | T-Value |
| --- | --- | --- |
| General Embryology | 0.21 | 2.08 |
| System-based embryology | 0.51 | 4.56 |
| Method of teaching | 0.24 | 2.15 |
| Educational assistance tools | 0.19 | 1.98 |

This study shows that the embryology course is effective on the clinical practice of medical students (P=0.04, CI=0.95). Descriptive statistics in general embryology showed that the topic of birth defects with the average score of 2.73 exerts the highest influence and the topic of the embryonic period with an average score of 2.32 exerts the lowest influence on the clinical practice of medical students.

### The effect of system-based embryology on the clinical performance of medical students

Standard pathway coefficient of system-based embryology with respect to the clinical practice of medical students was 0.51 (Table 2 and Figure 2). Thus, according to t value of this pathway (t=4.56 > 1.96) (Figure 1), it can be concluded (with 99% confidence) that system-based embryology affects the clinical performance of medical students. It can be estimated that 1 unit change in system-based embryology leads to 0.51 unit change in the clinical practice of medical students.

The topic of development of body cavities (diaphragm development) with an average score of 2.66 and the topic of eye and ear with an average score of 2.07 exert the highest and the lowest influence on the clinical practice of medical students, respectively.

### The effect of teaching method on the clinical performance of medical students

Table 2 shows that the standard pathway coefficient of the method of teaching with respect to the clinical performance of medical students was 0.24 (Figure 2). Thus, according to t value of this pathway (t=2.15 > 1.96) (Figure 1), it can be concluded (with 95% confidence) that the method of teaching affects the clinical practice of medical students. It can be estimated that 1 unit change in the method of teaching leads to 0.24 unit change in the clinical performance of medical students.

Lecturer’s ability to present the course with an average score of 3.40 and his/her seriousness in teaching with an average score of 3.13 has the highest and lowest effects on the clinical practice of medical students, respectively.

### The effect of educational assistance tools on the clinical performance of medical students

According to table 2, the standard pathway coefficient of the educational assistance tools with respect to the clinical practice of medical students was 0.19 (Figure 2). Therefore, according to t value of this pathway (t=1.98 > 1.96) (Figure 1), it can be concluded (with 95% confidence) that educational assistance tools affect the clinical performance of medical students. It can be estimated that 1 unit change in educational assistance tools lead to 0.19 unit change in the clinical performance of medical students.

Data showed that power point and video projector (with an average score of 3.22) is more effective than translated references (with an average score of 2.95).

## Discussion

Since embryology is of particular importance for medical students, and with respect to the fact that trainees and interns can challenge their knowledge of medicine in clinical performance, this study was conducted to elucidate the viewpoint of medical students (trainees and interns) of Zanjan University of Medical Sciences in Iran on the application of embryology course in their clinical performance. Briefly, our study indicates that the topic of birth defects is the most influential part of general embryology for clinical practice. In system-based embryology development of body cavities and diaphragm is of higher clinical importance from the viewpoint of trainees and interns in comparison to other topics. PowerPoint and video projector, and lecturers’ ability to present the course are other most important factors in the clinical practice of medical students.

Hamilton, *et al*. showed that general and system-based embryology courses are effective on the clinical practice of medical students [8]. Our results are in agreement with their reports. In addition, our study showed the importance of lecturers’ ability to present the course which is in line with the study of Kaimkhani, *et al*. who stated that lecturers’ capability in presenting the course and students’ preparation before the course is effective factors on embryology learning [9].

Wilson, *et al.* reported that educational assistance tools such as translated reference books, and PowerPoints presented by video projector has the most and least positive effects on anatomy learning, respectively [10]. It is in contrast to our results which show the importance of PowerPoint presented by video projectors. The difference may be the result of how well PowerPoints are designed and presented by lecturers.

Scott, *et al*. reported that embryology learning is of substantial importance in diagnosing and differentiating diseases according to the relative symptoms [4]. Our study demonstrates that embryology is most effective in finding a better approach to cure the disease and less effective on diagnosis from symptoms.

According to Carlson, Bruce. M [5], there are several methods of teaching in embryology: self-contained course – instructor independently determines contents and continuity, distributed model – essential elements are presented in a few lectures, system model – adopted by several medical books, problem-based method – exposure to embryology occurs in the content of the clinical problem. Finding the best approach to teach embryology which meets all the needs of medical students seems so difficult. With the decreasing volume of time allocated to anatomy and embryology teaching, undergraduate students must acquire a core understanding of anatomy, rather than knowing every detail of the human body [11]. Several studies have presented various approaches in presenting embryology including online courses with recorded lectures providing the ability to pause, replay and view any time and any place [12, 13], or the team-based learning (TBL) in which working with other students provides an effective way to learn content and promote clinical reasoning skills [14]. Emphasis on clinical problems in teaching embryology has been reported by several studies [6, 7] stating that clinically oriented embryology courses motivate students to consider embryology as an essential part of their medical career.

The present study indicated that a clinically oriented course may be more effective for clinical performance of clinical medical students. More emphasis on topics of system-based embryology assists medical students in their clinical practice. Educational assistance tools like 3-D models of the embryo and enhancing the quality of presentation of embryology courses are recommended.

## Acknowledgements

We would like to thank all the medical students of Zanjan University of Medical Sciences in pre-clinic and clinic phases for their excellent collaboration.

## Authors’ contributions

S.M. developed the concept and collected data and revised the manuscript, H.G. analysed data and revised the manuscript, M.J.M. collected data and revised the manuscript, A.F. developed the concept and collected data and revised the manuscript, M.G. collected data and drafted the manuscript, M.A. supervised the study and interpreted data and revised the manuscript critically. All authors read and approved the final manuscript.

## References

1. Sadler TWTW. Langman’s medical embryology. 12th ed: Lippincott Williams & Wilkins, a Wolters Kluwer business; 2012. 374 p.

2. Beale EG, Tarwater PM, Lee VH. A retrospective look at replacing face-to-face embryology instruction with online lectures in a human anatomy course. Anat Sci Educ. 2014;7:234–41.

3. Pourghasem M, Sum s. Practical Anatomy as an Advance Organizer for Anatomy Lectures: Effectiveness in Learning Facilitation for Dental Students. Iranian Journal of Medical Education. 2011;11:478–84.

4. Scott KM, Charles AR, Holland AJA. Clinical embryology teaching: is it relevant anymore? ANZ Journal of Surgery. 2013;83:709–12.

5. Carlson BM. Embryology in the medical curriculum. The Anatomical Record. 2002;269:89–98.

6. Moxham BJ, Emmanouil-Nikoloussi E, Standley H, Brenner E, Plaisant O, Brichova H, et al. The attitudes of medical students in Europe toward the clinical importance of embryology. Clinical Anatomy. 2016;29:144–50.

7. Zaletel I, Marić G, Gazibara T, Rakocevic J, Labudovic Borovic M, Puškaš N, et al. Relevance and attitudes toward histology and embryology course through the eyes of freshmen and senior medical students: Experience from Serbia. Annals of Anatomy - Anatomischer Anzeiger. 2016;208:217–21.

8. Hamilton J, Carachi R. Clinical embryology: is there still a place in medical schools today? Scott Med J. 2014;59:188–92.

9. Kaimkhani ZA AM, Fayez MA, Khoshhal K, Zafar M, Javaid A. Does the existing traditional undergraduate Anatomy curriculum satisfy the senior medical students. South East Asian Journal of Medical Education. 2010;3:20–6.

10. Wilson DR, Nava PB. Medical student responses to clinical procedure teaching in the anatomy lab. Clin Teach. 2010;7:14–8.

11. Mitchell R, Batty L. Undergraduate perspectives on the teaching and learning of anatomy*. ANZ Journal of Surgery. 2009;79:118–21.

12. Anstine J, Skidmore M. A Small Sample Study of Traditional and Online Courses with Sample Selection Adjustment. The Journal of Economic Education. 2005;36:107–27.

13. Aragon SR, Johnson SD, Shaik N. The Influence of Learning Style Preferences on Student Success in Online Versus Face-to-Face Environments. American Journal of Distance Education. 2002;16:227–43.

14. Nieder GL, Parmelee DX, Stolfi A, Hudes PD. Team-based learning in a medical gross anatomy and embryology course. Clin Anat. 2005;18:56–63.

